# High resolution spatial investigation of intracellular oxygen in muscle cells

**DOI:** 10.1101/2023.07.18.548845

**Authors:** Rozhin Penjweini, Alessandra Pasut, Branden Roarke, Greg Alspaugh, Dan L. Sackett, Jay R. Knutson

## Abstract

Molecular oxygen (O_2_) is one of the most functionally relevant metabolites. O_2_ is essential for mitochondrial aerobic respiration. Changes in O_2_ affect muscle metabolism and play a critical role in the maintenance of skeletal muscle mass, with lack of sufficient O_2_ resulting in detrimental loss of muscle mass and function. How exactly O_2_ is used by muscle cells is less known, mainly due to the lack of tools to address O_2_ dynamics at the cellular level. Here we discuss a new imaging method for the real time quantification of intracellular O_2_ in muscle cells based on a genetically encoded O_2_-responsive sensor, Myoglobin-mCherry. We show that we can spatially resolve and quantify intracellular O_2_ concentration in single muscle cells and that the spatiotemporal O_2_ gradient measured by the sensor is linked to, and reflects, functional metabolic changes occurring during the process of muscle differentiation.

**Highlights:**

- Real time quantitation of intracellular oxygen with spatial resolution
- Identification of metabolically active sites in single cells
- Oxygen metabolism is linked to muscle differentiation

## 1. Introduction

Molecular oxygen (O_2_) is one of the most functionally relevant metabolites. O_2_ is essential for mitochondrial aerobic respiration and O_2_ availability plays a central role in the cellular metabolism and energy production of many cells, including muscle cells, through oxidative phosphorylation (OXPHOS) [1, 2]. O_2_ also serves as a key metabolite in many other cytosolic and nuclear biochemical reactions implicated in gene/protein regulation [2].

Methods to trace O_2_ concentration ([O_2_]) or oxygen partial pressure (pO_2_) at the cell and mitochondrial level are highly desirable to fully understand the role of O_2_ in physiological as well as pathological conditions. However, the direct assessment of O_2_ dynamics at the intracellular level remains technically challenging, and most techniques for the study of O_2_ metabolism are either limited to bulk analysis (which do not resolve inter/intra cell heterogeneity) or they capture only time-integrated changes which do not reveal dynamic changes in [O_2_]. Hence, despite its importance, how exactly O_2_ is used by muscle cells to regulate metabolism has been given little attention. To address this knowledge gap, the intracellular pO_2_ level was monitored in single muscle cells at different stages during differentiation via the use of our recently developed (FRET)-based O_2_ sensor [3, 4]. Further, we also exploit the autofluorescence of metabolic co-factors nicotinamide adenine dinucleotide (NAD(P)H) and flavin adenine dinucleotide (FAD) to reveal changes in cell metabolism. We show that muscle cells are primarily oxidative in nature but retain the ability to use glycolysis when cultured in low glucose medium and in their undifferentiated state. Further, we unexpectedly observed substantial heterogeneity in intracellular oxygenation, and we showed how changes in the media-imposed pO_2_ (from 1 to 136 mmHg) influence intracellular and mitochondrial pO_2_ distributions as cells transit from a single to a multinucleated state. We provide evidence that mitochondrial O_2_ consumption and NAD(P)H oxidases are linked to and reflect functional metabolic changes occurring during the process of muscle differentiation.

## 2. Materials and Methods

### 2.1. Design of the O_2_ probe

The O_2_ sensitive probe pMyo-mCherry was generated as previously described [3, 4]. Briefly, pMyo-mCherry was prepared by cloning the myoglobin gene from Physeter catodon (Sperm Whale, Addgene Plasmid pMB413a, #20058) into the pmCherry N1 vector (Clonetech, Mountain View, CA). A 2-amino acid (Ser-Gly) spacer was inserted between the C-terminus of myoglobin and the N-terminus of mCherry to allow for flexibility and avoid possible misfolding [5]. A mitochondrial targeting sequence from the mouse mitochondrial transcription factor A (TFAM) was added to the pMyo-mCherry vector to generate a mitochondrial (mtMyo-mCherry) O_2_ sensor [4]. The detailed description of the probe mechanism of action can be found elsewhere [3].

### 2.2. Cell transfection with O_2_ probe

C2C12 cells, a mouse skeletal muscle cell line, were kept in Dulbecco’s modified Eagle’s medium (DMEM, Gibco, Grand Island, NY, USA) supplemented with 10% fetal bovine serum, and 1% peni-cillin-streptomycin solution (Mediatech Inc. Manassas, VA, USA). The cells were plated in μ-Slide 4 well or 8 well chambers (Ibidi GmbH, Martinsried, Germany) at a density of 2.0-2.5 × 10^4^ cells/cm^2^. Transfection was performed in antibiotic free medium using FuGENE^®^ HD transfection reagent (Promega Corporation, Durham, NC, USA) following the manufacturer’s instructions with a final plasmid amount of ∼100-200 ng per well. After 24 h, the transfection media was removed, and the cells were incubated in fresh culture medium for an additional 24 h before imaging. To induce muscle differentiation, C2C12 cells were grown to 70-80% confluency before switching to differentiation medium (5% horse serum in DMEM). Cells were kept in differentiation medium for up to 6 days with alternate days feeding. For glucose starvation studies, transfected C2C12 cells (both proliferative and differentiated) were switched to no-glucose, no glutamine medium for 24 h before imaging. After live imaging, cells were fixed in 4% PFA, permeabilized in Triton 0.05%, and immunoassayed for tubulin (anti-α-tubulin (clone DM1A), 1:100 dilution in blocking buffer) to detect cell morphology and en-dogenous mCheery fluorescence.

### 2.3. Treatment with 2,4-Dinitrophenol (DNP) and Rotenone/Antimycin A

Transfected C2C12 cells grown in monolayer were kept in cell media with 50 μM DNP (mitochondrial uncoupler) or a mixture of 1 μM rotenone/antimycin during the imaging. DNP transports protons across the mitochondrial inner membrane, altering the proton gradient and inhibiting ATP production via OXPHOS. Rotenone inhibits the transfer of electrons from complex I to co-enzyme Q (CoQ), whereas antimycin prevents the oxidation of CoQ by cytochrome *c*.

### 2.4. Imaging set up

Two-photon FLIM was performed using a Leica SP5 confocal laser scanning microscope (Buffalo Grove, IL) equipped with a tunable Chameleon Ti:Sapphire femtosecond laser (Coherent, UK) operating at 80 MHz with wavelengths set to 720, 780, and 880 nm for the excitation of NAD(P)H, Myo-mCherry and FAD, respectively. The laser light was passed through a 685 nm LP dichroic mirror and directed to a Leica Plan-Apochromat 40×, 1.1 N.A. water immersion microscope objective (laser power ≤ 7 mW at the objective). A 680 nm short pass filter and a 560 nm long pass dichroic mirror were used to respectively reduce the laser scattering and to split the NAD(P)H and FAD fluorescence toward two hybrid photomultiplier detectors (HyD, Leica Microsystems). 460/60 nm, 552/57 nm and 647/57 nm bandpass filters (Semrock BrightLine^®^, Rochester, NY, USA) were used to further filter NAD(P)H, FAD and mCherry signals, respectively. The electrical pulse output from the HyD was directed into an SPC-150 photon counting card (Becker & Hickl, Berlin, Germany). Synchronization with the pixel, line, and frame clock from the scanning unit of the microscope was used for image construction in time-correlated single photon counting (TCSPC) mode. Single cells were imaged for 50-80 s (depending on the intensity), image size was set to 256 × 256 (pixels)^2^, and TCSPC histograms were collected with 256 channels in a 12.5 ns time window.

### 2.5. Controlled imposed pO_2_ during imaging

A miniature incubation chamber with a gas mixing system (CO_2_−O_2_−MI, Bioscience Tools, San Diego, CA, USA) was mounted onto the microscope stage to keep the temperature at 37 °C and provide 5% CO_2_ and a stable %O_2_ (v/v) of 18.5%, 10%, 5% or 0.5% during the imaging; 0.5% is the lowest %O_2_ attainable by our system. The cells in culture dishes with a ∼ 3 mm layer of medium above them, without lids reached a stable pO_2_ within 45 min when monitored by a 250 μm diameter bare-fiber O_2_ sensor (NX-BF/O/E, Optronix Ltd., Oxford, UK) connected to an OxyLite, 1 Channel monitor (Optronix Ltd., Oxford, UK). The media-imposed external pO_2_ (in mmHg) was measured at the bottom of the 4-well or 8-well chambers with or without live cells present. The Oxylite fiber sensor has a 250um hemispherical tip, so this imposed level is actually the level in media averaged from ∼0-150um above the bottom.

### 2.6. Intracellular mapping of the pO_2_, free/bound NAD(P)H and FLIRR

The activity of OXPHOS and glycolytic metabolism was monitored in proliferative and differentiated C2C12 cells by lifetime imaging of Myo-mCherry and metabolic co-factors NAD(P)H and FAD at different imposed pO_2_. The fluorescence lifetime decays of Myo-mCherry, NAD(P)H and FAD were obtained by a double-exponential decay model in SPCImage (Becker & Hickl GmbH, Berlin, Germany) at optimized goodness of fit (*χ*^2^). As described previously [3, 4], the decay curves at each pixel were fit using a non-linear least-squares method to follow:

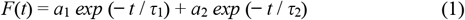

where *a*_1_ and *a*_2_ are pre-exponential factors and can be used (if natural lifetime is constant) to represent the fraction of fluorophores with shorter (*τ*_1_) and longer (*τ*_2_) lifetimes, respectively.

The parameters *a*_1_% and *a*_2_% for NAD(P)H and FAD were generated via amplitude weighting for each pixel to calculate the free/bound NAD(P)H (*a*_1,NAD(P)H_%/*a*_2,NAD(P)H_%) [6-9] and fluorescence life-time based redox ratio (FLIRR; *a*_2,NAD(P)H_%/*a*_1,FAD_%) [10].

For the pO_2_ measurements, the mean lifetime of Myo-mCherry was obtained for each single image and averaged across multiple cells (n > 30). Then, using a method that was described previously [3, 4], the resulting lifetime values (*τ*(pO_2_)) were plotted against the media-imposed external pO_2_ and a hyperbolic curve was fit to the data using the Curve Fitting Toolbox in MATLAB R2020b (The Math-Works Inc., Natick, Massachusetts):

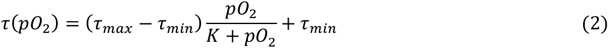

where *τ*_max_ and *τ*_min_ are the longest and shortest average lifetime for Myo-mCherry at normoxia and hypoxia, respectively. *K* is a fitting parameter related to the affinity of myoglobin for O_2_.

To obtain intracellular pO_2_, the *τ*(pO_2_) of actively respiring C2C12 cells were compared to those treated with rotenone/antimycin (incapable of significant O_2_ consumption). The intracellular pO_2_ in rotenone/antimycin treated cells is assumed to be equivalent to the media-imposed pO_2_. Therefore, the *τ*(pO_2_) values for the treated cells can be used as a reference for the lifetime of the probe at the environmental level of pO_2_ present in solution. Rearranging Eq. (2), it is possible to back calculate the effective pO_2_ at each lifetime value, fixing the *K* and *τ*_*max*_ to the values obtained from the rotenone/antimycin data.

Finally, pseudo color mapping of pO_2_, free/bound NAD(P)H and FLIRR in the intracellular environment was obtained using MATLAB R2019b (The MathWorks Inc.) equipped with the Image Processing Toolbox. More detailed description can be found somewhere else [3].

### 2.7. Statistical Analyses

For each condition, FLIM was recorded in at least 30 cells. Kruskal-Wallis and Mann–Whitney U tests were used to evaluate whether the values in the independent groups are significantly different from each other. Analyses were carried out using SPSS 14.0 (a subsidiary of IBM, Chicago, IL, USA) software and statistical significance was defined at *p* < 0.05 (95% confidence level).

## 3. Results and Discussions

### 3.1. Building spatial mapping of intracellular pO_2_ at the single cell level

To enable the quantification of intracellular O_2_ with spatiotemporal resolution at the single cell level, C2C12 cells, a murine muscle cell line, were transfected with Myo-mCherry sensor (Fig 1A). Myo-mCherry is a Förster resonance energy transfer (FRET)-based probe formed by Myoglobin (O_2_-binding protein) and the fluorophore mCherry, linked together. The main principle of this sensor is that the binding of O_2_ to Myoglobin results in changes in the FRET-depleted emission intensity of mCherry. These changes in mCherry intensity can be detected by fluorescence lifetime imaging such that when the probe is deoxygenated (low [O_2_]) its emission intensity and correspondingly its lifetime will decrease compared to its oxygenated form (high [O_2_]) (Fig 1B).

**Figure 1.**
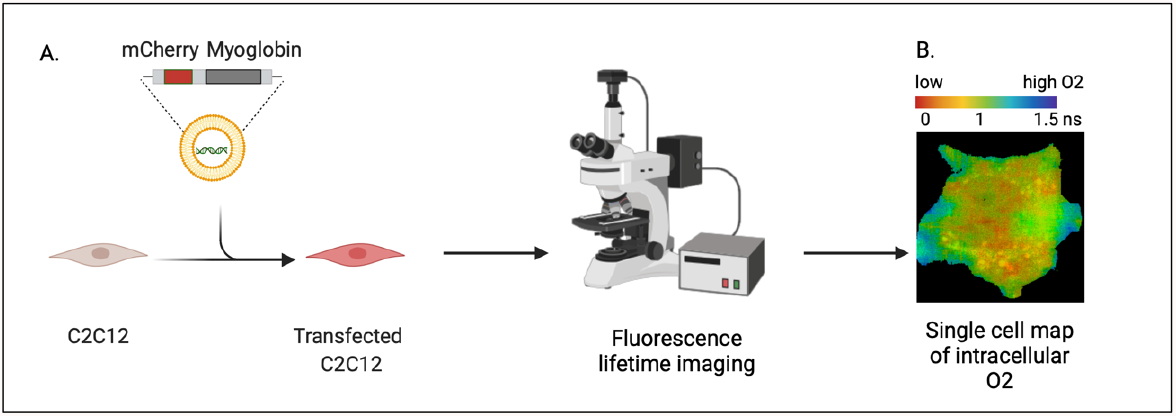
Schematic of the set-up for the spatial map of intracellular [O_2_] in single cells. A. To monitor intracellular pO_2_, the C2C12 murine cell line was transiently transfected with the Myo-mCherry plasmid and FLIM recording acquired after 48 hours from the transfection. B. Representative pseudo color image of a transfected C2C12 cell. Color reflects changes in lifetime (measured in ns) and used as a proxy of intracellular O_2_ concentration.

To better resolve intracellular oxygenation (and explain functionally relevant regional differences in pO_2_), we used two different versions of the Myo-mCherry sensor: one targeting the whole cell and one specific to the mitochondria. We confirmed the correct localization of the probes via immunofluorescence of the mCherry signal (Supp Fig 1A). Further, both probes were well tolerated over long period of time by both proliferating cells (kept in growth medium in high serum conditions) and post-mitotic multinucleated cells (switched to low serum medium) with no obvious effect on cell behavior or differentiation potential of the cells (Supp. Fig 1 and Suppl. Fig 2).

### 3.2. Quantitative assessment of intracellular -derived pO_2_ during muscle differentiation

First, changes in Myo-mCherry lifetime in response to different imposed pO_2_ (from ∼136 to ∼1 mmHg, corresponding to a range from 18.5% to 1% oxygen) were measured in the whole cell and in mitochondria of the proliferative and differentiated C2C12 cells.

As shown in Figs. 2 and 3, at all imposed pO_2_ levels, Myoglobin-mCherry had shorter lifetime values in the mitochondria of both proliferative (see Fig. 2A) and differentiated cells (myotubes with two or more nuclei) compared to whole cell values (cytoplasm) (see Fig. 3A). These data indicate that mitochondria are the primary sites for of O2 consumption in muscle cells and therefore can be distinguished for having lower [O2] compared to other intracellular regions. In addition, we found that differentiated muscle cells consume more O2 compared to undifferentiated cells [11].

**Figure 2.**
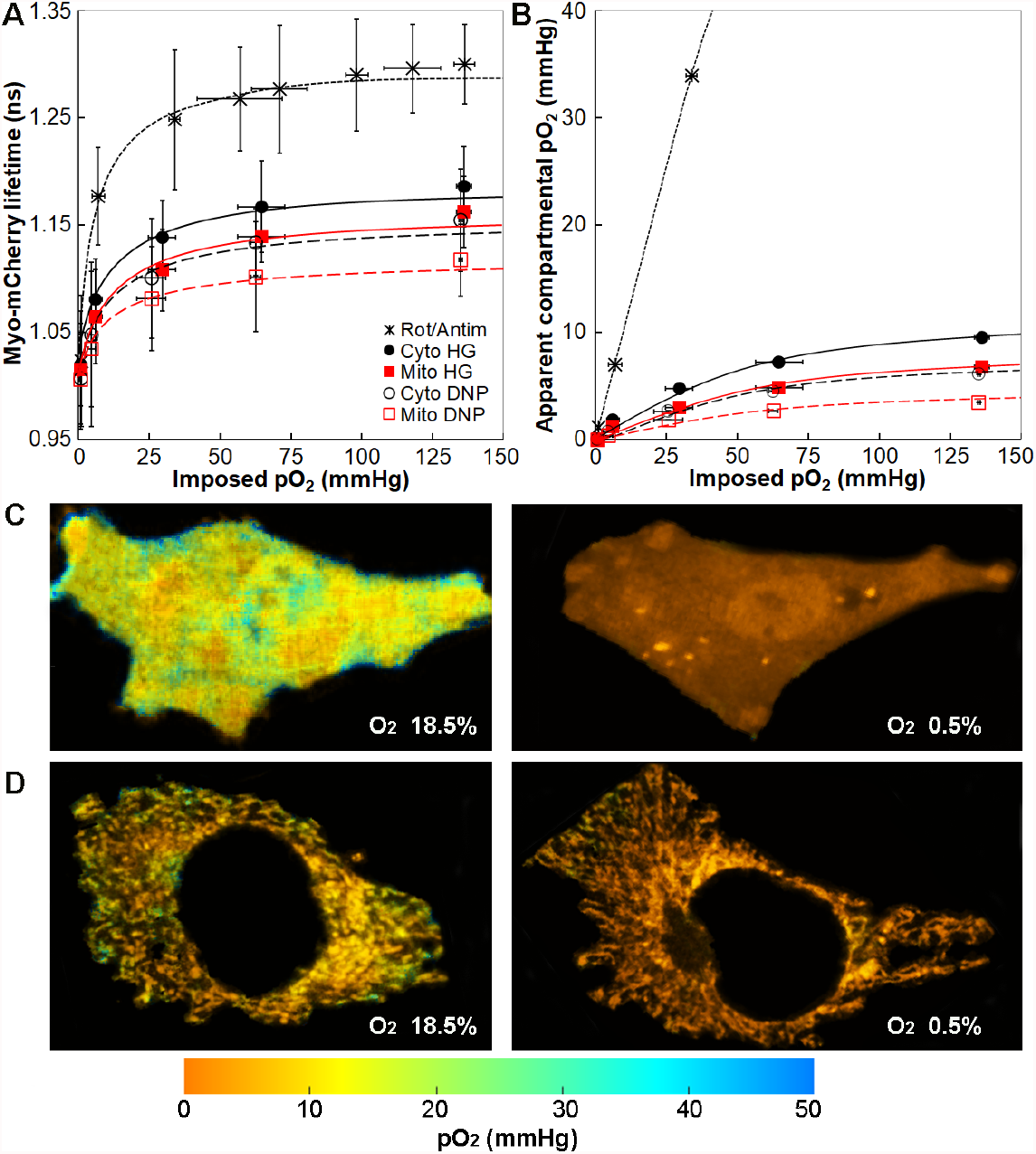
Intracellular pO_2_ level obtained from Myo-mCherry lifetime in proliferative C2C12 cells. A) Average fluorescence lifetime of Myo-mCherry within cell (black filled circles) and mitochondria (red filled rectangles) versus imposed pO_2_. The lifetime values are compared to those in the cells treated with rotenone/antimycin (black crosses) or DNP (empty black circles and red rectangles). The data are shown with the best hyperbolic fit obtained from Eq. (2). Vertical and horizontal error bars are the standard deviations obtained from at least 30 cells in three independent experiments. B) The apparent intracellular pO_2_ calculated from the FLIM data in proliferative cells by using the rotenone/antimycin calibration curve in A see also Methods) are plotted versus the media-imposed pO_2_. Pseudocolor mapping of apparent C) cytosolic and D) mitochondrial pO_2_ in the intracellular environment of proliferative cells in response to imposed O_2_% of 18.5 (left side) and 0.5% (right side). In the color bars, red indicates lower pO_2_ values, whereas blue indicates higher values. Intracellular pO_2_ distribution was obtained from the Myo-mCherry lifetime images and rotenone/antimycin calibration curve.

**Figure 3.**
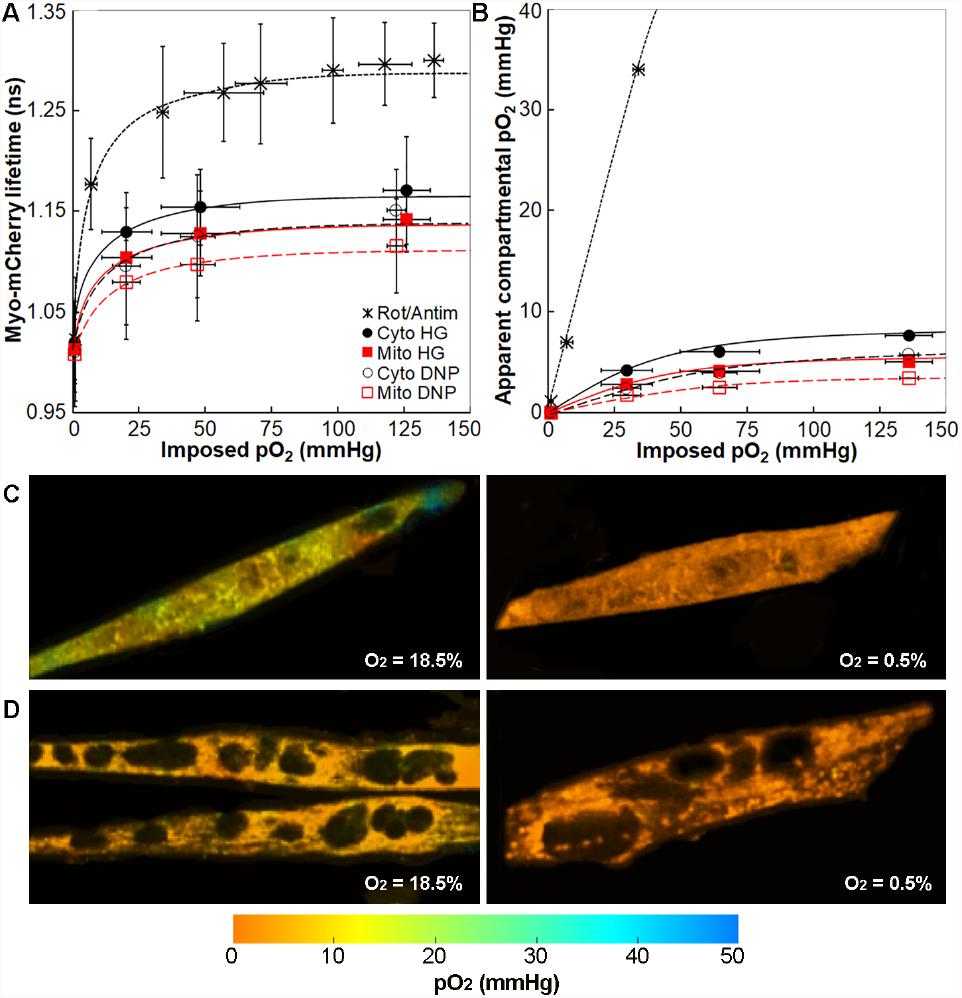
Intracellular pO_2_ level obtained from Myo-mCherry lifetime in differentiated C2C12 cells. A) Average fluorescence lifetime of Myo-mCherry in cytosol (black filled circles) and mitochondria (red filled rectangles) of C2C12 cells versus the imposed pO_2_. The lifetime values are compared to those in the cells treated with rotenone/antimycin (black crosses) or DNP (empty black circles and red rectangles). The data are shown with the best hyperbolic fit obtained from Eq. (2). Vertical and horizontal error bars are the standard deviations obtained from at least 30 cells in three independent experiments. B) The apparent intracellular pO_2_ calculated from the FLIM data in proliferative cells by using the rotenone/antimycin calibration curve in A (see also Methods) are plotted versus the media-imposed pO_2_. Pseudo-color mapping of apparent C) cytosolic and D) mitochondrial pO_2_ in the intracellular environment of proliferative cells in response to imposed O_2_% of 18.5 (left side) and 0.5% (right side). In the color bars, red indicates lower pO_2_ values, whereas blue indicates higher values. Intracellular pO_2_ distribution was obtained from the Myo-mCherry lifetime images and rotenone/antimycin calibration curve.

Next, we challenged C2C12 with different mitochondria stressors (a mixture of rotenone/antimycin inhibitors and DNP uncoupler). As expected, the lifetime of Myo-mCherry increased in both the cytoplasm and mitochondria of the cells treated with short pulse mixture of rotenone/antimycin (inhibitors of mitochondrial OXPHOS). Of note, the treatment did not affect cell survival. The increase in lifetime values was comparable between undifferentiated and differentiated cells (∼9% in cytosol and ∼11% in mitochondria of undifferentiated C2C12 cells and ∼10% in cytosol and ∼12% in mitochondria of differentiated cells when compared to the untreated cells at the highest imposed pO_2_).

On the contrary, in the presence of the DNP uncoupler, the average lifetime values of Myo-mCherry were shorter in both the cytoplasm and mitochondria (as shown in Figs. 2A and 3A). The lifetime of Myo-mCherry in DNP-treated undifferentiated C2C12 decreased by ∼3% in cytosol and ∼4% in mitochondria at the highest imposed pO_2_ when compared to the untreated cells. In differentiated C2C12 treated with DNP, the lifetime values decreased by ∼2% in the cytosol and mitochondria at the highest imposed pO_2_ when compared to the untreated cells. As shorter lifetime indicates a lower pO_2_ [3], these data show that in the presence of DNP, C2C12 cells increase their oxidative metabolism and consume O_2_ at their highest rate in both proliferative and differentiated states. The increase of O_2_ consumption after the addition of DNP is reported as being a result of dissociation between oxidative and phosphorylative processes [12].

Finally, we sought to quantify intracellular pO_2_ distribution. As reported previously [3], the life-time of Myo-mCherry as a function of pO_2_ follows the hyperbolic O_2_ dissociation behavior of myoglobin. Therefore, the best hyperbolic fit to the lifetime data versus imposed pO_2_ was obtained using Eq. (2) (fitting parameters available in Table S1) and the intracellular versus imposed pO_2_ was then obtained (Figs. 2B and 3B). Specifically, the rotenone/antimycin fit curve was used to calibrate the Myo-mCherry sensor *in situ*, assuming mitochondrial O_2_ consumption in C2C12 cells is turned off by these stressors. Mapping via the rotenone/antimycin fit curve formula in Eq. (2) allows one to transform and replot Figs. 2A and 3A as intracellular versus imposed pO_2_ as seen in Figs. 2B and 3B.

From the graphs shown Figs. 2A and 3A we conclude, in agreement with the results obtained from lifetime values, that (i) mitochondria consume more oxygen than other regions of the cell as evidenced by the fact that mitochondrial values of pO_2_ are lower than “cytoplasmic” (all interior) values of pO_2_ at all imposed pO_2_; (ii) the sustained internal pO_2_ levels are well below those applied revealing the O_2_ consumption in these cells. Note that we did not assume the media-imposed pO_2_ is identical to that found at the cell surface to recover the apparent internal pO_2_; that is only strictly needed for the reference curve with rotenone/antimycin.

These results are also supported by pseudocolor mapping of the pO_2_ distribution as shown in Figs. 2C-D and 3C-D where a shift toward red indicates lower pO_2_ values, and blue indicates higher pO_2_ values. Of note, pO_2_ distribution is not homogenous within individual cells with perinuclear regions having lower pO_2_ values compared to the cell edges (Figs 2C-D).

### 3.3. pO_2_ correlates with cell metabolic states

In our previous result, we showed that the intracellular pO_2_ in individual muscle cells can be used to infer the metabolic state of the cells and that differentiated muscle cells consume more oxygen than undifferentiated cells. To independently confirm such findings and to monitor the activity of OXPHOS and glycolytic metabolism in proliferative and differentiated C2C12 cells, lifetime imaging of the metabolic co-factors NAD(P)H and FAD was performed at different imposed O_2_%. At high pO_2_, NAD(P)H is converted to NAD^+^ by the enzyme NAD(P)H dehydrogenase [13] and NAD(P)H exists predominantly in an enzyme bound state (defined by lower *a*_1_%/ *a*_2_% where *a*_*1*_ and *a*_*2*_ represent the two bound and unbound components), whereas FADH_2_ is converted to non-enzyme bound FAD by the enzyme succinate dehydrogenase (lower bound FAD fraction, thus higher *a*_2_%/*a*_1_%) [10]. Free and bound populations of NAD(P)H as well as fluorescence lifetime redox ratio (FLIRR) were correlated to the corresponding imposed O_2_%.

FLIM results indicate that differentiated muscle cell have lower glycolysis and more active mitochondria capable of higher O_2_ consumption compared to their proliferative counterpart, as evidenced by higher FLIRR (Figs. 4A and C) and lower free NAD(P)H (Supp. Fig. 3A and C). In hypoxic condition, FLIRR was higher at higher pO_2,_ and it decreased by ∼6% in proliferative cells and by ∼7% in differentiated cells upon hypoxia (see Figs. 4A and C). Of note, pseudo-color mapping of free/bound NAD(P)H ratio (*a*_1_%/*a*_2_%) and FLIRR revealed a substantial degree of metabolic heterogeneity in both proliferative (Fig. 4B) and differentiated cells (Fig. 4D) possibly indicating mitochondrial bioenergetic heterogeneity.

**Figure 4.**
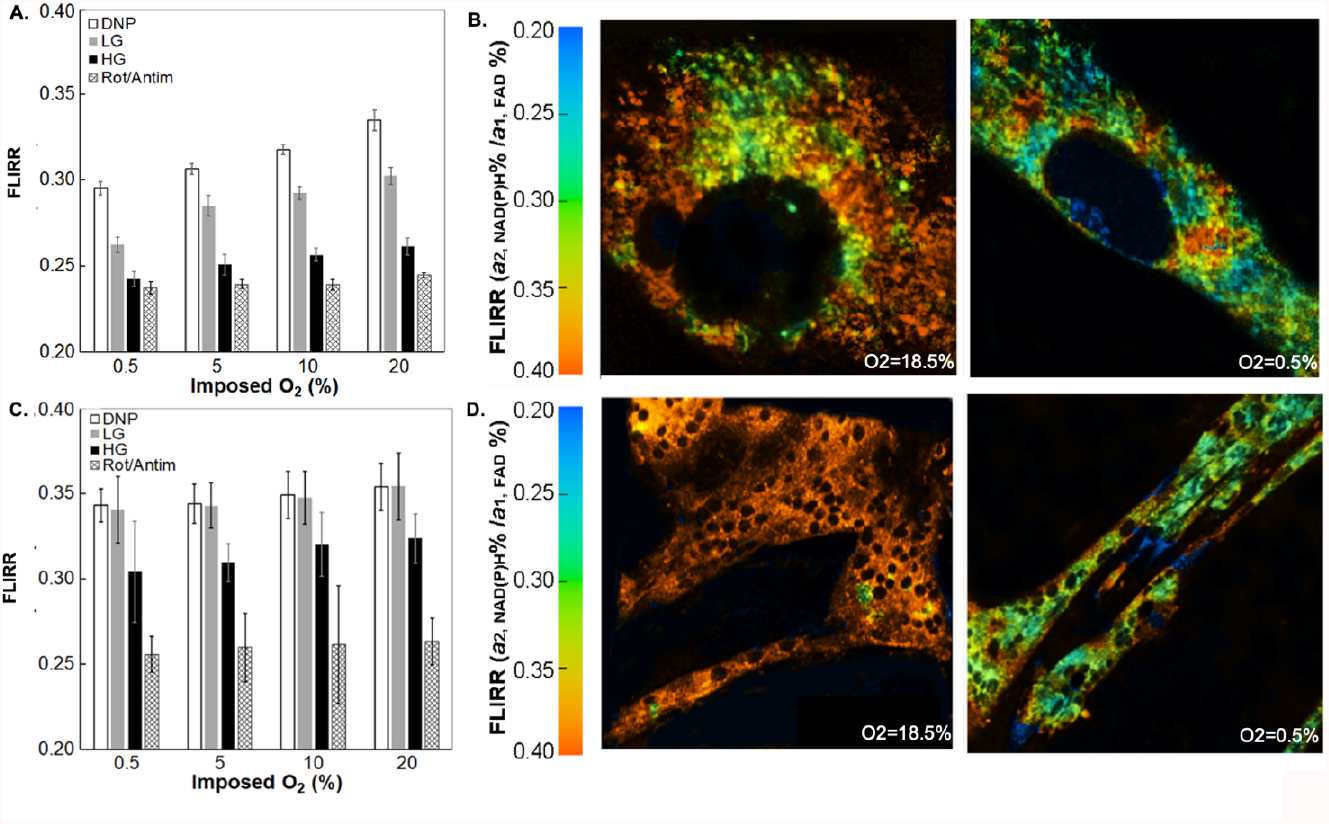
Changes in fluorescence lifetime redox ratio (FLIRR) in proliferative (A-B) and differentiating (C-D) C2C12 cells. Bar graphs show changes in FLIRR (*a*_2,NAD(P)H_%/*a*_1, FAD_%) versus imposed O_2_ (%) for pro-liferative (A) and differentiating (C) C2C12 across 4 tested conditions. The error bars are the standard deviations from at least 30 cells. Pseudo color mapping of FLIRR in the intracellular environment of proliferative (B) and differentiating (D) cells in response to imposed O_2_ of 18.5% and 0.5%. In the color bars, red indicates lower values, whereas blue indicates higher values of FLIRR.

The effect of O_2_ consumption on free and bound NAD(P)H production and FLIRR was further studied by disrupting O_2_ consumption with rotenone/antimycin and enhancing O_2_ consumption with DNP or low glucose media. As shown in Figs 4A and C, lower FLIRR was observed in rotenone/antimycin treated conditions indicating a switch from OXPHOS to glycolysis in both differentiated and proliferative cells. In agreement, higher free NAD(P)H production was noted in rotenone/antimycin treated conditions (Supp Fig 3A and C). Similar results were observed when cells were cultured in low glucose media (more physiological condition) (Figs 4A and C; Suppl Fig 3 A and C).

## 4. Conclusions

Molecular oxygen (O_2_) is a key substrate for mitochondrial ATP production with changes in oxygen affecting cell metabolism and ultimately playing a critical role in organ development but also pathological states (ischemia, cancer) [14]. Further, metabolic heterogeneity is recognized as a key feature of cell plasticity [15] and genetically encoded metabolite sensors are particularly useful to study intracellular metabolite dynamics and metabolic adaptations at the single cell level.

Here we described a FLIM-based method to spatially resolve and quantify intracellular oxygenation and metabolic state in muscle cells in two different cell states (proliferative and post-mitotic), all at subcellular resolution. We used two different oxygen probes (whole cell and mitochondria targeted) and we showed that differentiated muscle cells are oxidative and use glycolysis when cultured in hypoxic conditions/low glucose or in their undifferentiated state. We show evidence that mitochondria are the primary sites of oxygen consumption. We unexpectedly also captured a substantial degree of metabolic heterogeneity at the mitochondrial level which requires further investigation. While in this study we applied the sensor to the investigation of mitochondrial O_2_ metabolism, we envision that nuclear, membrane or peroxisome-targeted oxygen sensors will be useful to also address non-mitochondrial sources of O_2_ consumption.

## Supplementary materials

Supplementary materials is provided that includes: Organelle-targeted sensor; long-term culture assessment; Changes of free and enzyme-bound NAD(P)H in proliferative and differentiated C2C12 cells; OxyLite measurements of the imposed pO_2_; Parameters of the hyperbolic fits;

## Author Contributions

R.P. and A.P.; Designed the experiments, prepared and transfected the cells, performed various cell treatments and wrote the paper. A.P.; Examined the Myo-mCherry transfection efficiency and cytotoxicity. R.P. and B.R.; Performed OxyLite measurements, FLIM data collection and analyses. R.P. and G.A.; Performed fluorescence confocal imaging, cell treatment and data analysis for detection of the carbonyl groups and mitochondrial ROS. G.A.; Generated pseudo color mapping of pO_2_ and FLIRR. D.L.S. and J.R.K.; Supervised the research.

## Funding and Acknowledgments

This work was supported by the Intramural Research Program of the National Heart, Lung, and Blood Institute (NHLBI), and the Eunice Kennedy Shriver National Institute of Child Health and Human Development (NICHD), National Institutes of Health (NIH). Alessandra Pasut was supported by funding from the NIH Office of Intramural Training and Education (OITE). We would like to acknowledge the Light Microscopy Core at NHLBI for the use of their confocal microscopes for fluorescence lifetime imaging and Dr. Evelyn Ralston and Aster Kenea for initial support and for providing C2C12 cells.

