## Supplementary files for "High resolution spatial investigation of intracellular oxygen in muscle cells"

**Supplementary Figures**

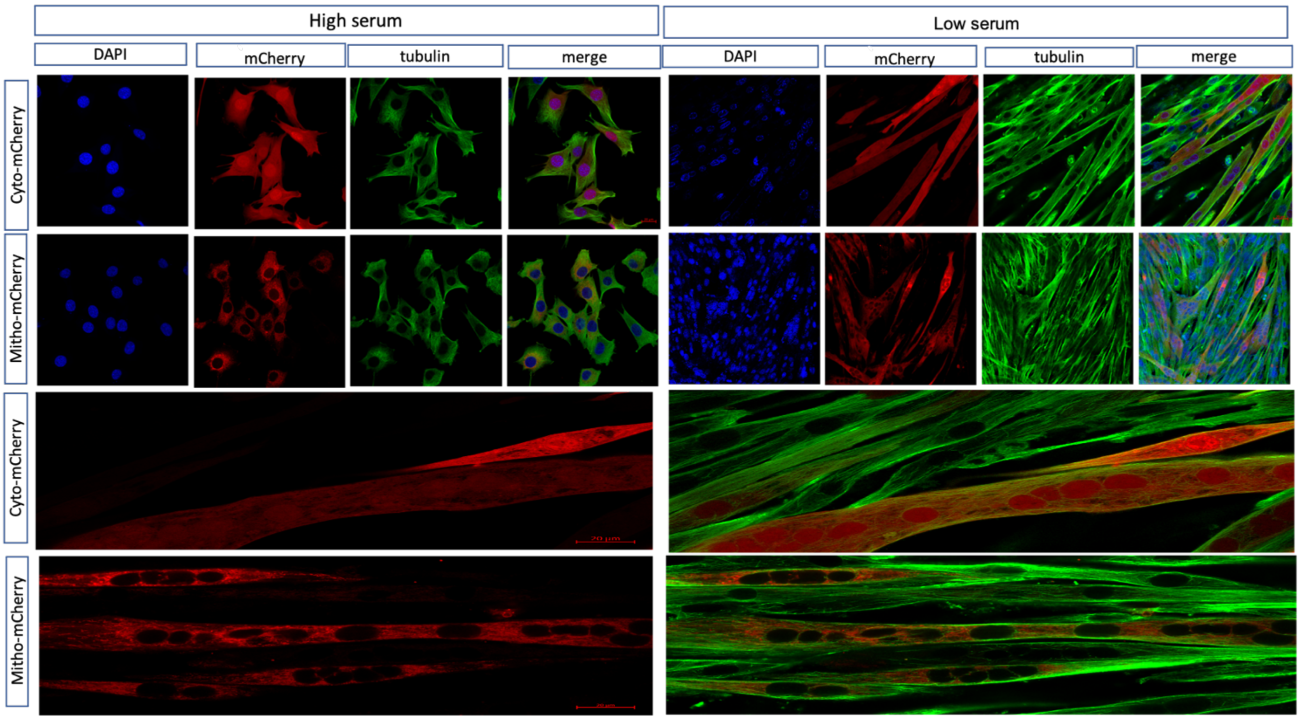


**Supp Fig 1. Organelle-targeted sensor.** Upper panels. Representative immunofluorescence of C2C12 cells transiently transfected with cytoplasmic/whole cell (cyto-mCherry) and mitochondrial (mito-mCherry) oxygen probes and grown in proliferative (high serum) and differentiating (low serum) conditions. Lower panel: high magnification of fully differentiating C2C12 cells. mCherry shown in red; tubulin shown in green, nuclei counterstained with DAPI shown in blue.

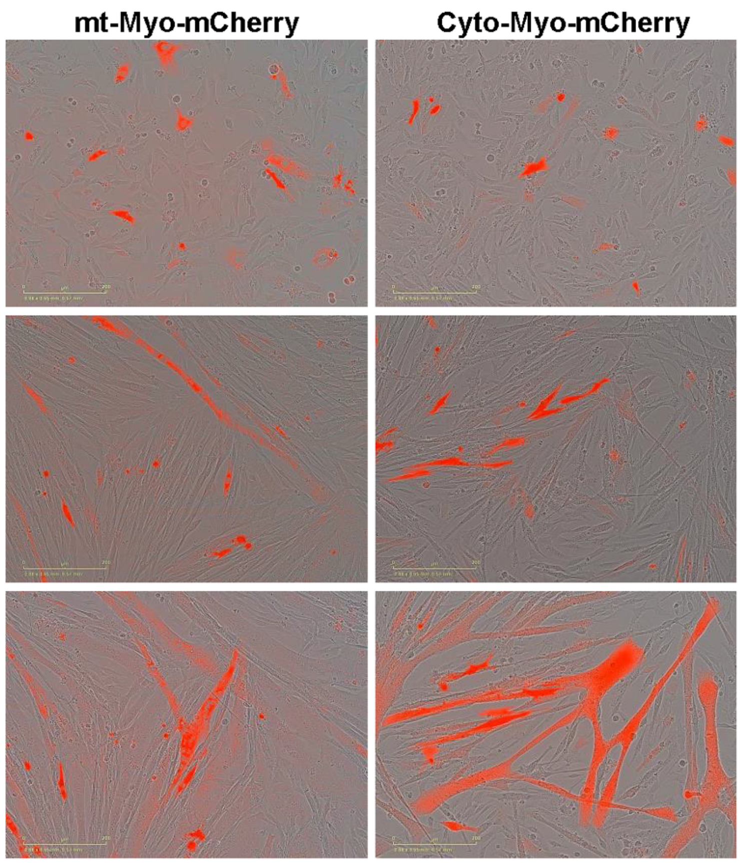


**Supp Fig 2. Long term assessment.** C2C12 cells were transfected with either cytoplasmic/whole cell (left) or mitochondrial (right) targeted sensor and cultured in high serum and differentiation media. Upon switching to differentiation media, elongated myotubes expressing mCherry fluorescence could be detected in the transfected cells for up to 5 days.

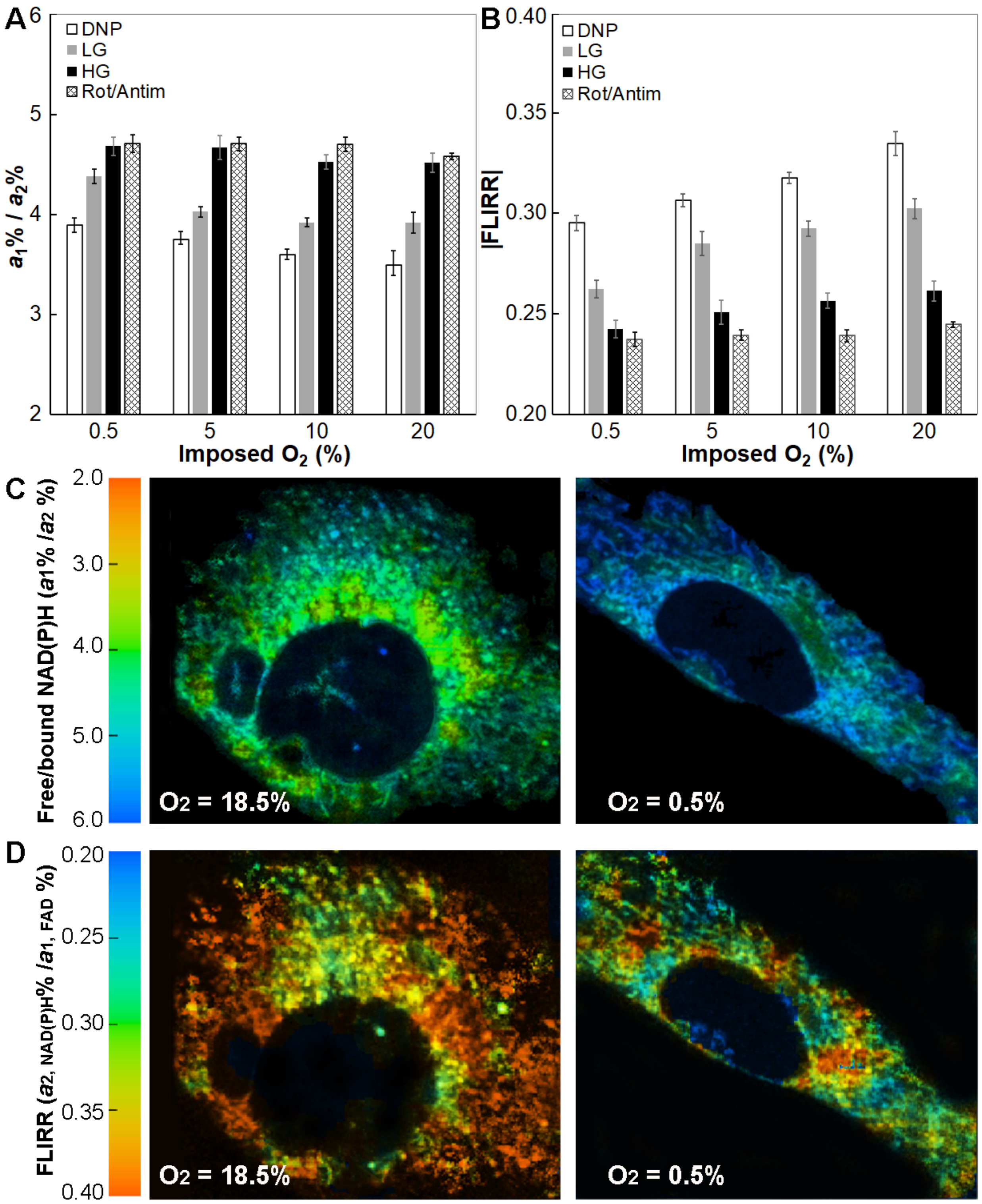


**Supp Fig 3. Changes of free and enzyme-bound NAD(P)H in proliferative and differentiated C2C12 cells.** Bar graphs show changes in free/bound NAD(P)H ratio ($a_1\%/a_2\%$), versus imposed $O_2\%$ for proliferative (A) and differentiating (C) C2C12 across 4 tested conditions. The error bars are the standard deviations from at least 30 cells. Pseudocolor mapping of free and enzyme-bound NAD(P)H ratio ($a_1\%/a_2\%$) in the intracellular environment of proliferative (B) and differentiating (D) cells in response to imposed $O_2$ of 18.5% and 0.5%. In the color bars, red indicates lower values, whereas blue indicates higher values of $a_1\%/a_2\%$.

**Table S1.** OxyLite measurements of the $pO_2$ level at the C2C12 cell layer (proliferative or differentiated in the bottom of petridish) treated with stressors DNP or rotenone/antimycin at different imposed $O_2$%. The data are shown with standard deviation.

| | **Proliferative C2C12** | | | **Differentiated C2C12** | | |
|---|---|---|---|---|---|---|
| **$O_2$ (%)** | **$pO_{2,\ Control}$ (mmHg)** | **$pO_{2,\ DNP}$ (mmHg)** | **$pO_{2,\ Rot/Antim}$ (mmHg)** | **$pO_{2,\ Control}$ (mmHg)** | **$pO_{2,\ DNP}$ (mmHg)** | **$pO_{2,\ Rot/Antim}$ (mmHg)** |
| 0.5 | 0.93 ± 0.12 | 0.80 ± 0.20 | 1.93 ± 0.60 | 0.80 ± 0.17 | 0.80 ± 0.20 | 2.83 ± 0.60 |
| 5 | 29.56 ± 4.85 | 25.78 ± 5.12 | 33.92 ± 1.96 | 20.37 ± 9.49 | 20.23 ± 5.12 | 33.92 ± 1.96 |
| 10 | 64.55 ± 8.30 | 62.61 ± 1.65 | 71.01 ± 16.00 | 48.17 ± 14.91 | 47.10 ± 6.50 | 76.01 ± 16.00 |
| 18.5 | 135.92 ± 2.62 | 134.87 ± 0.49 | 136.25 ± 6.00 | 126.00 ± 8.87 | 122.00 ± 3.50 | 136.25 ± 6.00 |

$pO_{2,\ control}$ are for the cells with no stressor

**Table S2.** Parameters of the hyperbolic fits. Fitting parameter *K* was obtained from fitting the data presented in Figs. 2-3A to Eq. (2). $\tau_{max}$ is the longest average lifetime for Myo-mCherry in each cell type measured at normoxia ($O_2$ = 18.5%). Each parameter is shown with its standard deviation.

| | **Undifferentiated C2C12 cells** | | | | **Differentiated C2C12 cells** | | | |
|---|---|---|---|---|---|---|---|---|
| | **$\tau_{min}$ (ns)** | **$\tau_{max}$ (ns)** | ***K*** | **$R^2$** | **$\tau_{min}$ (ns)** | **$\tau_{max}$ (ns)** | ***K*** | **$R^2$** |
| Cytosol | 1.02 ± 0.04 | 1.18 ± 0.04 | 11.25 ± 5.17 | 0.98 | 1.02 ± 0.04 | 1.17 ± 0.05 | 7.99 ± 6.31 | 0.98 |
| Mitochondria | 1.01 ± 0.05 | 1.16 ± 0.03 | 12.54 ± 5.93 | 0.98 | 1.01 ± 0.03 | 1.14 ± 0.03 | 8.11 ± 7.47 | 0.97 |
| Cyto/DNP | 1.01 ± 0.02 | 1.15 ± 0.05 | 12.53 ± 9.21 | 0.98 | 1.01 ± 0.04 | 1.15 ± 0.04 | 10.08 ± 7.45 | 0.98 |
| Mito/DNP | 1.01 ± 0.04 | 1.12 ± 0.03 | 13.68 ± 9.51 | 0.98 | 1.01 ± 0.05 | 1.12 ± 0.05 | 10.89 ± 7.41 | 0.98 |
| Mito/Rot/Antim | 1.02 ± 0.06 | 1.30 ± 0.04 | 6.55 ± 2.98 | 0.96 | 1.02 ± 0.01 | 1.30 ± 0.02 | 6.55 ± 2.98 | 0.96 |
